# Biogeographical and ecological factors associated with kill rates of an apex predator

**DOI:** 10.1101/2020.10.04.325779

**Authors:** Bogdan Cristescu, L. Mark Elbroch, Justin A. Dellinger, Wesley Binder, Christopher C. Wilmers, Heiko U. Wittmer

## Abstract

Kill rates and functional responses are fundamental to the study of predator ecology and the understanding of predatory-prey dynamics. As the most widely distributed apex predator in the western hemisphere pumas (*Puma concolor*) have been widely studied yet a biogeographical synthesis of their kill rates is currently lacking. We reviewed the literature and compiled data on sex- and age-specific kill rate estimates of pumas on ungulates, and conducted analyses aimed at understanding ecological factors explaining the observed variation across their range. Kill rate studies on pumas, while numerous, were primarily conducted in Temperate Conifer Forests (< 10% of puma range), revealing a dearth of knowledge across much of their range, especially from tropical and subtropical habitats. Across studies, kill rates in ungulates/week were highest for adult females with kitten(s) (1.24 ± 0.41 ungulates/week) but did not vary significantly between adult males (0.84 ± 0.18) and solitary adult females (0.99 ± 0.26). Kill rates in kg/day did not differ significantly among reproductive classes. Kill rates of adult pumas increased with ungulate density. Ungulate species richness had a weak negative association with adult male kill rates. Neither scavenger richness, the proportion of non-ungulate prey in the diet, nor regional human population density had a significant effect on ungulate kill rates. Our results had a strong temperate-ecosystem bias highlighting the need for further research across the diverse biomes pumas occupy in order to make species level inferences. Data from more populations would also allow for multivariate analyses providing deeper inference into the ecological and behavioural factors driving kill rates and functional responses of pumas, and apex predators in general.

## Introduction

Kill rates, defined as the number of prey or biomass killed by an individual predator per unit time, are of continued interest to ecologists and wildlife managers. A predator’s functional response describes how kill rates vary with prey density (Holling 1959) and is fundamental in predicting the stability threshold for prey populations under the impacts of predation, as well as in estimating the potential carrying capacity of predator populations (Carbone and Gittleman 2002; Sinclair 2003; Dunn and Hovel 2020). Both kill rates and functional responses, however, are influenced by diverse ecological variables and are thus difficult to extrapolate beyond local scales (Zimmermann et al. 2015).

Estimating kill rates for most carnivores is logistically challenging, expensive and time consuming. For these reasons, biogeographical meta-analyses of large carnivore kill rates that may offer insights into the ecological variables driving them have rarely been accomplished. This is a concern given that kill rate estimates are needed to further theoretical modeling of functional responses and predator-prey dynamics, and to develop effective conservation strategies for predators and prey in a changing world (e.g. the effects of climate change on functional response to predict future predator-prey dynamics and stability; Rall et al. 2012).

Estimates of kill rates alone, however, are inadequate to determine the effects of predation on prey populations (Vucetich et al. 2011). For example, kill rates do nothing to elucidate whether predation is additive or compensatory, which has real implications for determining the need for managing predators and their prey. Only when kill rates are combined with information on predator density data can they be scaled up to estimates of “total predation rates”, and, when used in conjunction with additional knowledge including vital rates and population size of the prey, inform management actions in complex multi-species systems (Owen-Smith and Mills 2008; Bonenfant et al. 2009; Forrester and Wittmer 2013). Understanding the link between predation and prey density is particularly important for small prey populations declining due to apparent competition (e.g. Johnson et al. 2013).

Because carnivore kill rates are difficult to obtain, most studies on carnivore foraging instead focus on prey composition (i.e. frequency of occurrence), and in some cases biomass of prey species in a carnivore’s diet. Many carnivores consume a wide range of prey, which evidence suggests is due to variable prey availability (Hayward et al. 2006; Hayward et al. 2016), prey catchability/accessibility (Hopcraft et al. 2005; Balme et al. 2007), and the age or life history stage of the carnivore itself (Hayward et al. 2007; Elbroch et al. 2017a; Blecha et al. 2018; Elbroch and Quigley 2019). When used in conjunction with prey availability, diet composition data enable researchers to estimate prey selection, or preference (Hayward et al. 2006). Prey preference for rare prey, for example, is suggestive of the potential negative impacts predators may have on rare prey population dynamics (Elbroch and Wittmer 2013a), but like kill rates alone, is inadequate to determine the effects of predation on the respective prey.

Nevertheless, kill rate studies have been recently conducted for large carnivores tagged with VHF or GPS collars (e.g. gray wolves, *Canis lupus*, Sand et al. 2005; jaguars, *Panthera onca*, Cavalcanti and Gese 2010; tigers, *Panthera tigris*, Miller et al. 2013; leopards, *Panthera pardus*, Farhadinia et al. 2018). However, few studies have tried to explain what may be driving large carnivore kill rates beyond explanatory factors at local scale (but see Elbroch et al. 2014). The lack of data on carnivore kill rates and the inconsistency with which they are studied remains pervasive, precluding a more holistic understanding of carnivore foraging ecology across biomes. This is the case even for widely studied carnivores such as pumas (*Puma concolor*), which to our knowledge, were the first apex predator for which kill rate estimation was attempted (Connolly 1949).

Although pumas consume a variety of prey (Martínez-Gutiérrez et al. 2015), ungulates comprise a large proportion of their diet across their range (Murphy and Ruth 2009). Pumas are ideally sized to capture and subdue deer (*Odocoileus spp.;* Carbone et al. 1999), their primary prey throughout much of North America (Murphy and Ruth 2009). Nonetheless, pumas in general hunt the most common ungulate species, which in some regions of North America is elk (*Cervus elaphus*) (e.g. Elbroch et al. 2013), and in parts of South America is the vicuña (*Vicugna vicugna*) (Smith et al. 2019) or guanaco (*Lama guanicoe*) (Elbroch and Wittmer 2013a). Pumas can regularly kill prey as large as moose (*Alces alces*) and feral horses (*Equus caballus*) (Knopff et al. 2010).

We compiled puma foraging studies from across their biogeographical distribution and extracted kill rate values from those papers that reported them. We also derived study area-specific ecological variables that might explain variation in puma kill rates across their range. The overall goal was to improve our understanding of predation and our ability to inform conservation management of both predator and prey species. We first assessed biogeographical variability in kill rates exhibited by different puma reproductive classes. We then contrasted factors that might influence localized kill rates and tested the following predictions:

1. Kill rates differ between females and males, and between solitary females and females with dependent young (Laundré 2005), because of differences in energetic needs among reproductive classes.
2. Prey availability, specifically density and ungulate species richness, positively correlates with kill rates (i.e. in the case of prey density, a Type I functional response; Holling 1959).
3. The abundance of alternative prey (e.g. non-ungulate species) inversely correlates with kill rate (i.e. prey switching for an abundant alternative prey reduces predation on primary prey; Elbroch et al. 2015a; Keehner et al. 2015; Soria-Díaz et al. 2018).
4. Puma kill rates positively correlate with species richness of scavengers that are dominant to pumas, as kleptoparasitism drives pumas to kill additional prey (Krofel et al. 2012; Elbroch and Wittmer 2013b; Elbroch et al. 2015b).
5. Kill rates positively correlate with human density, since pumas are fearful of people (Smith et al. 2017) and exhibit reduced handling times having to kill more frequently near people (Smith et al. 2015).
6. Field methodology employed to study pumas impacts estimates of kill rates. Specifically, we predicted higher kill rate values for studies utilizing Global Positioning System (GPS) compared with those using Very High Frequency (VHF) collars (Merrill et al. 2010).

## Materials And Methods

### Literature Search

Our review of the literature on puma kill rates included North and South America. We carried out a search in Google Scholar in October 2019, using the keywords “puma”, “mountain lion”, “cougar”, or “*Puma concolor*” in conjunction with “kill rate”, “inter-kill interval”, or “diet”. We chose Google Scholar because it retrieves more records than alternative online databases such as Web of Science or Scopus (Harzing and Alakangas 2016). This was important because we wanted to maximize the coverage of graduate theses, book chapters and scientific reports, to supplement peer-reviewed journal publications. Some researchers who investigated puma kill rate in their study systems also summarized kill rates in tabular format for previous work (e.g. Anderson and Lindzey 2003; Knopff et al. 2010). We perused references listed in the respective tables and added them to our review if we had not identified them in the Google Scholar searches. We inspected puma management plans for state jurisdictions in the western U.S. and Canada to extract potential additional references, focusing on the sections that presented puma diet and ungulate relationships. We further augmented the list of studies by compiling relevant publications among our co-authors’ collections, and also contacted study authors to request digital copies of publications when needed. In all cases, when the same study was included in a graduate thesis as well as a peer-reviewed article, we retained the article for analysis. Only studies that monitored > 1 individual puma were considered.

### Study Areas

Once kill rate studies were identified, we delineated polygons corresponding to study area boundaries by digitizing their perimeters in Google Earth. A small subset of publications did not provide a study area figure. For these studies we generated the polygons based on figures available in other publications by the same research group, or on information in the “Study area” section, which described geographic and/or management landmarks. We exported the resulting polygons as vector shapefiles for use in GIS.

### Biogeography of Research Effort

We assessed the biogeographical research effort on puma kill rates to identify research gaps, by inspecting the distribution of studies according to major habitat types (i.e. biomes) defined by the World Wildlife Fund (*sensu* Olson and Dinerstein 1998), and publicly available from The Nature Conservancy (http://maps.tnc.org/gis_data.html). We converted the biome polygons to a raster file with a 1 km^2^ resolution, which has been used previously for puma spatial research at broad scale (Teichman et al. 2013). We clipped the raster to match current puma distribution as mapped by the International Union for Conservation of Nature (Nielsen et al. 2015). We employed the raster analysis toolbox to summarize raster values for each biome within puma range, and then calculated the percentages of biomes for the global puma distribution. We used Q-GIS v3.10.3. for all GIS procedures.

### Kill Rate Values

We extracted kill rate values as ratio estimators in number of ungulates (ungulates/week), and ungulate biomass killed per unit time (kg/day). Some studies reported kill rate as an inter-kill interval (days/kill), which we converted to ungulates/week. When seasonal kill rates were presented, we averaged them to obtain annual estimates. Studies in which livestock accounted for ≥ 5% of puma diet (*n* = 2) were excluded. We used a subset of studies that reported kill rate by puma reproductive class to inspect variability in kill rate among puma sex and age classes. We performed Kruskal-Wallis rank sum tests for kill rate by reproductive class pooled across biomes, as well as by biome if sample sizes allowed. We then carried out pairwise-comparisons between reproductive classes, using a Bonferroni adjustment for multiple comparisons (Dunn 1961).

### Factors Associated with Kill Rate

We used univariate linear regressions to investigate factors that may be associated with annual puma kill rates on ungulates. We tested five predictors: prey availability, puma diet, scavenger diversity, human disturbance, and field method for kill rate estimation (Table 2). For continuous covariates (Table 2) we also ran univariate generalized additive models (GAM) to investigate potential non-linear relationships with kill rate. Each GAM fitted a penalized regression spline with automated selection of knots for the spline. We used restricted maximum likelihood (REML) as smoothness selection criterion, because it is less sensitive to small sample sizes than other criteria (Wood 2011).

We carried out separate analyses for each adult reproductive class, but subadults were excluded due to small sample sizes. We modelled factors potentially associated with kill rate only for studies that involved locating kills based on GPS or VHF tracking of collared pumas (Supplementary Data S1). These studies either relied on field visitation to locate kills and calculate empirical kill rates or used data from confirmed kills in a predictive modelling framework. The latter studies typically applied location cluster algorithms parameterized with data from known kill sites to predict kill rates. We did not include studies based on energetic modelling, due to differences in their kill rate estimation as compared to intensive field studies (Knopff et al. 2010; Elbroch et al. 2014). For multiple-ungulate systems, we excluded studies that only calculated kill rate for one ungulate species of management interest (*n* = 2).

We calculated several measures of prey availability, including density, and richness (number of species) estimated in various manners (Supplementary Data S2, S3). We obtained ungulate density (ungulates/km^2^) directly from the puma kill rate studies that reported it, or by searching the literature for ungulate studies in the same region. In many cases, density estimates were extracted from state agency reports. Ungulate density data were summed across ungulate species that occurred ≥ 5% in puma diet in the respective study area, to obtain an overall ungulate density (*UngDens*) variable. We removed studies for which we were unable to find density of ungulates contributing ≥ 5% to diet. We derived several measures of ungulate species richness, including overall richness (*UngRichAll*); richness of ungulate species that occurred ≥ 5% in puma diet (*UngRichMain*); and richness of large-bodied ungulates (> 200 kg; elk, moose, feral horse) that contributed ≥ 5% to puma diet (*UngRichLarge*).

Pumas in populations that commonly feed on non-ungulate prey (*NonUng*) might have lower ungulate kill rates. Therefore, we considered a *NonUng* variable that encompassed the summed proportions of non-ungulate prey items in the diet of pumas for each system. Livestock were included as non-ungulate prey in calculation of the *NonUng* variable because they were not wild ungulates. The proportions were calculated based on the diet composition of pumas reported in the kill rate study, or in some cases in other publications for the respective study area. We used puma diet data from carcasses identified at GPS clusters, or via VHF monitoring of collared pumas, because scat studies would have likely overestimated the proportion of small items in puma diet (Bacon et al. 2011).

We considered scavengers and humans as potential sources of external disturbance that could influence puma kill rate. We calculated *ScavRich* as the richness of scavengers present in the study area. Only dominant facultative scavengers that can displace pumas from their kills were included in calculations. These include grizzly bear (*Ursus arctos*), American black bear (*U. americanus*), gray wolf, and jaguar (Elbroch and Kusler 2018). To calculate scavenger richness, we perused the study area description of kill rate studies for listing of the 4 species above. In the rare cases when this information was absent, we inspected government agency reports on dominant predator species distribution (WDFW 2008; Wiles et al. 2011).

We used *HumDens* to denote the density of humans in the study region (inhabitants/km^2^). We used census density for the respective county, or averaged densities across counties if the study area polygon spanned > 1 county (USA: www.census.gov, https://usa.ipums.org; Canada: www12.statcan.gc.ca). Density data were available per mile^2^, which we then converted to values per km^2^. Human population censuses did not occur every year, therefore we selected the census year that was closest to the middle year of the respective kill rate study.

The method used to identify predation events by collared predators could affect estimation of the kill rate value. Specifically, VHF techniques could possibly underrepresent predation compared to GPS cluster technology, by missing kills. We therefore generated *Method* as a categorical variable (1 = GPS, 2 = VHF) to test for possible effects of methodology on kill rate estimates.

Exploratory analysis showed that the kill rate models (ungulates/week) had improved fit when *UngDens* and *HumDens* were natural log-transformed, whereas models for biomass killed (kg/day) had best fit when the log transformation was applied to *UngDens* only. We therefore applied these transformations throughout the respective model sets. We ranked predictions based on ΔAICc and AICc weights, performing separate ranking for predictions tested with linear models (*n* = 8) and those tested with generalized additive models for continuous covariates (*n* = 3). We assessed model fit accounting for sample size with the adjusted coefficient of determination (*R*^2^). For supported models (ΔAICc < 2), we checked the linear regression assumptions including normality of residual distributions and homogeneity of error variances. We plotted residuals vs. fitted values, a normal Q-Q plot of standardized residuals, and a scale-location plot of square root of standardized residuals vs. fitted values.

Because sample sizes were relatively small, we assessed the robustness of regression outputs by investigating the effects of influential observations using Cook’s distance (*D*_*i*_), where 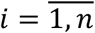. We considered observations to have high influence if 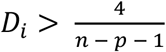 (Bruce and Bruce 2017), where *n* is the number of observations and *p* the number of predictor variables. We removed records with high influence from supported models and re-ran the models thereafter, taking note of the differences in outputs.

### Functional Response

To further investigate the relationship between prey density and kill rate, we generated functions for Type I, II and III functional responses (Holling 1959). We fitted the curves for each function and compared their performance by investigating the associated predictive errors. We calculated the root mean squared error (RMSE) for each model as an average measure of the deviations of the observed kill rate values from the fitted curves. Functional responses were investigated separately for each puma reproductive class.

We used program R v3.6.3 base functions in RStudio v1.1.447 as well as packages bbmle, ggplot2, mgcv and tidyverse for statistical analyses and for graphical outputs. We used Q-GIS v3.10.3 to generate a map illustrating the global distribution of studies on puma kill rate.

## Results

We reviewed 134 studies related to puma diet, of which 54 publications reported puma kill rate on ungulates (Supplementary Data S1). Most kill rate studies were graduate theses (*n* = 25) and peer-reviewed journal articles (*n* = 24), with many of the articles reporting on graduate research projects. Some articles reported data for > 1 study area. We found a small number of book chapters, symposium proceedings and published reports on the topic (*n* = 5). Once we removed duplicate studies, including graduate theses that were also published in journals, as well as studies that used kill rate information across multiple publications, the dataset included 30 study areas (Fig. 1).

**Fig. 1.**
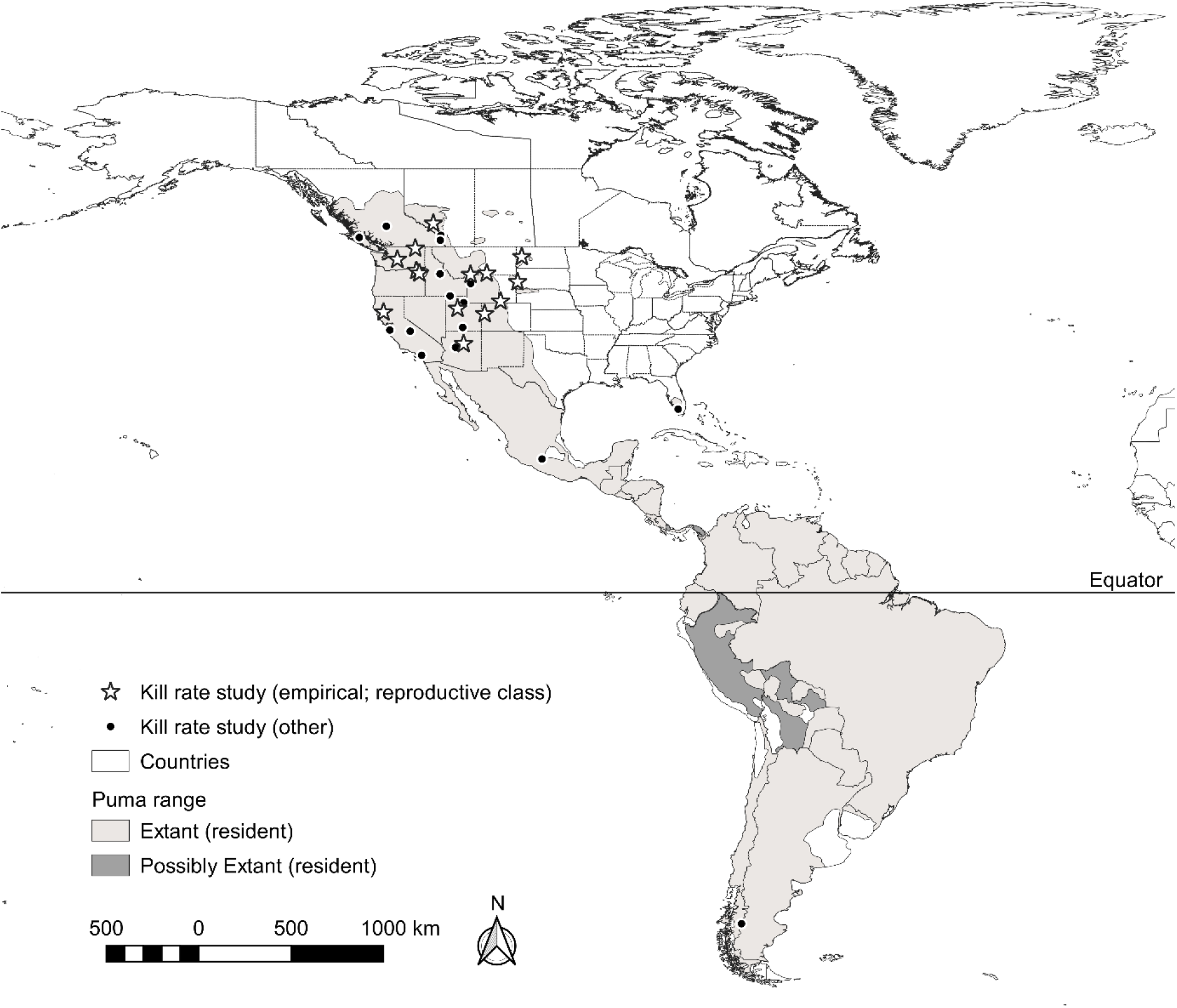
The geographical distribution of studies on puma kill rate overlaid with puma distribution range (Nielsen et al. 2015). Studies that we included in the analysis are illustrated with stars. Boundaries of states and provinces are illustrated for USA and Canada.

### Biogeography of Research Effort

Puma kill rate studies reporting by reproductive class (adult male, solitary adult female, adult female with kitten(s), subadult male, subadult female) disproportionately occurred in the northern hemisphere (Fig. 1). We were only able to find one study in the southern hemisphere (Elbroch et al. 2014), but the study was excluded from analysis due to high proportion of livestock in puma diet. All studies included in analyses were from North America, primarily from the USA.

Studies have disproportionally focused on Temperate Conifer Forests (< 10% of puma range) (Table 1), which hosted 11 of 14 studied reporting in ungulates/week, and 5 of 7 studies reporting in kg/day. One study each occurred in Desert and Xeric Shrublands; Temperate Grasslands, Savannas and Shrublands; and Boreal Forest/Taiga. We did not find reports on puma kill rate from kill site investigations in Tropical and Subtropical regions, which represent 36% of puma range (Table 1).

**Table 1.** Biomes in the global distribution range of the puma. Research effort on puma kill rate is expressed as number of studies by biome.

| Biome | Global<br>range<br>(%) | Kill rate<br>studies<br>(ungulates/<br>week) | Kill rate<br>studies<br>(kg/day) |
| --- | --- | --- | --- |
| Tropical and Subtropical Moist Broadleaf Forests | 36 | 0 | 0 |
| Deserts and Xeric Shrublands | 15 | 1 | 0 |
| Tropical and Subtropical Grasslands, Savannas and<br>Shrublands | 13 | 0 | 0 |
| Temperate Conifer Forests | 9 | 11 | 5 |
| Temperate Grasslands, Savannas and Shrublands | 8 | 1 | 1 |
| Tropical and Subtropical Dry Broadleaf Forests | 8 | 0 | 0 |
| Montane Grasslands and Shrublands | 3 | 0 | 0 |
| Mediterranean Forests, Woodlands and Scrub | 2 | 0 | 0 |
| Temperate Broadleaf and Mixed Forests | 2 | 0 | 0 |
| Tropical and Subtropical Coniferous Forests | 2 | 0 | 0 |
| Boreal Forests/Taiga | 1 | 1 | 1 |
| Flooded Grasslands and Savannas | 1 | 0 | 0 |
| Mangroves | <1 | 0 | 0 |

Inland water, as well as rock and ice (combined < 1% of puma range) were excluded because they were not puma habitat. Only studies that reported kill rate based on field investigations, by reproductive class and in systems where livestock did not contribute significantly ( $\geq 5\%$ ) to puma diet are listed. Studies had to report kill rate for at least 1 of 5 reproductive classes to
be included. Reproductive classes considered were adult male, solitary adult female, adult female with kitten(s), subadult male, and subadult female.

**Table 2.** Predictor variables used in modelling puma kill rate across its global distribution range.

| Predictor | Description | Code | Type | Units | Value<br>range<sup>1</sup> | Value<br>range<sup>2</sup> |
| --- | --- | --- | --- | --- | --- | --- |
| Prey<br>availability | Ungulate density ($\geq 5\%$ in puma diet) | *UngDens* | Continuous | Ungulates<br>/km<sup>2</sup> | 0.59 –<br>50.75 | 0.73 – 50.75 |
|  | Ungulate species richness (overall) | *UngRichAll* | Categorical | Species | 2 – 8 | 2 – 8 |
| | Ungulate species richness ($\geq 5\%$ in puma diet) | *UngRichMain* | Categorical | Species | 1 – 4 | 1 – 4 |
| | Large ungulate species richness ($\geq 5\%$ in puma diet) | *UngRichLarge* | Categorical | Species | 0 – 2 | 0 – 2 |
| Puma diet | Contribution of non-ungulate prey to puma diet | *NonUng* | Continuous | Percentage | 0 – 19 | 4.62 – 18.18 |
| Scavenger<br>diversity | Species richness of dominant scavengers | *ScavRich* | Categorical | Species | 0 – 3 | 0 – 3 |
| Human<br>disturbance | Human density | *HumDens* | Continuous | Inhabitants/<br>km<sup>2</sup> | 2.39 –<br>414.10 | 2.39 – 31.60 |
| Method | Technological approach to estimate kill rate | *Method* | Categorical | Unitless | 1 – 2 | NA (GPS<br>only) |

<sup>1</sup> Value range for modelling kill rate frequency (ungulates/week). <sup>2</sup> Value range for modelling biomass killed (kg/day). Only studies that reported kill rate based on field investigations, by reproductive class and in systems where livestock did not contribute significantly ( $\geq 5\%$ ) to
puma diet are listed. Studies had to report kill rate for at least 1 of 5 reproductive classes to be included. Reproductive classes considered were adult male, solitary adult female, adult female with kitten(s), subadult male, and subadult female.

### Kill Rate Values

#### Frequency

Kill rates (ungulates/week) among 14 studies varied by adult puma reproductive status (Kruskal-Wallis 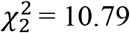, *P* = 0.005). Specifically, kill rates of adult males (mean ± *SD* ungulates/week, 0.84 ± 0.18) differed from those of adult females with kittens (1.24 ± 0.41) (pairwise comparison Wilcoxon rank sum with Bonferroni adjustment, *P* = 0.008) but not from solitary adult females (0.99 ± 0.26) (Fig. 2A). Kill rates of subadults (male 0.77 ± 0.13, female 0.80 ± 0.32) appeared lower than for adults, but sample sizes were too small for analysis. When subsampled by studies that occurred in Temperate Conifer Forests (*n* = 11), kill rate also differed by adult reproductive status (Kruskal-Wallis 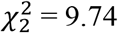, *P* = 0.008). Similar to the previous finding, kill rates differed between adult males and adult females with kittens (pairwise comparison Wilcoxon rank sum with Bonferroni adjustment, *P* = 0.019), but also between solitary females and females with kittens (*P* = 0.077, *α* = 0.10).

**Fig. 2.**
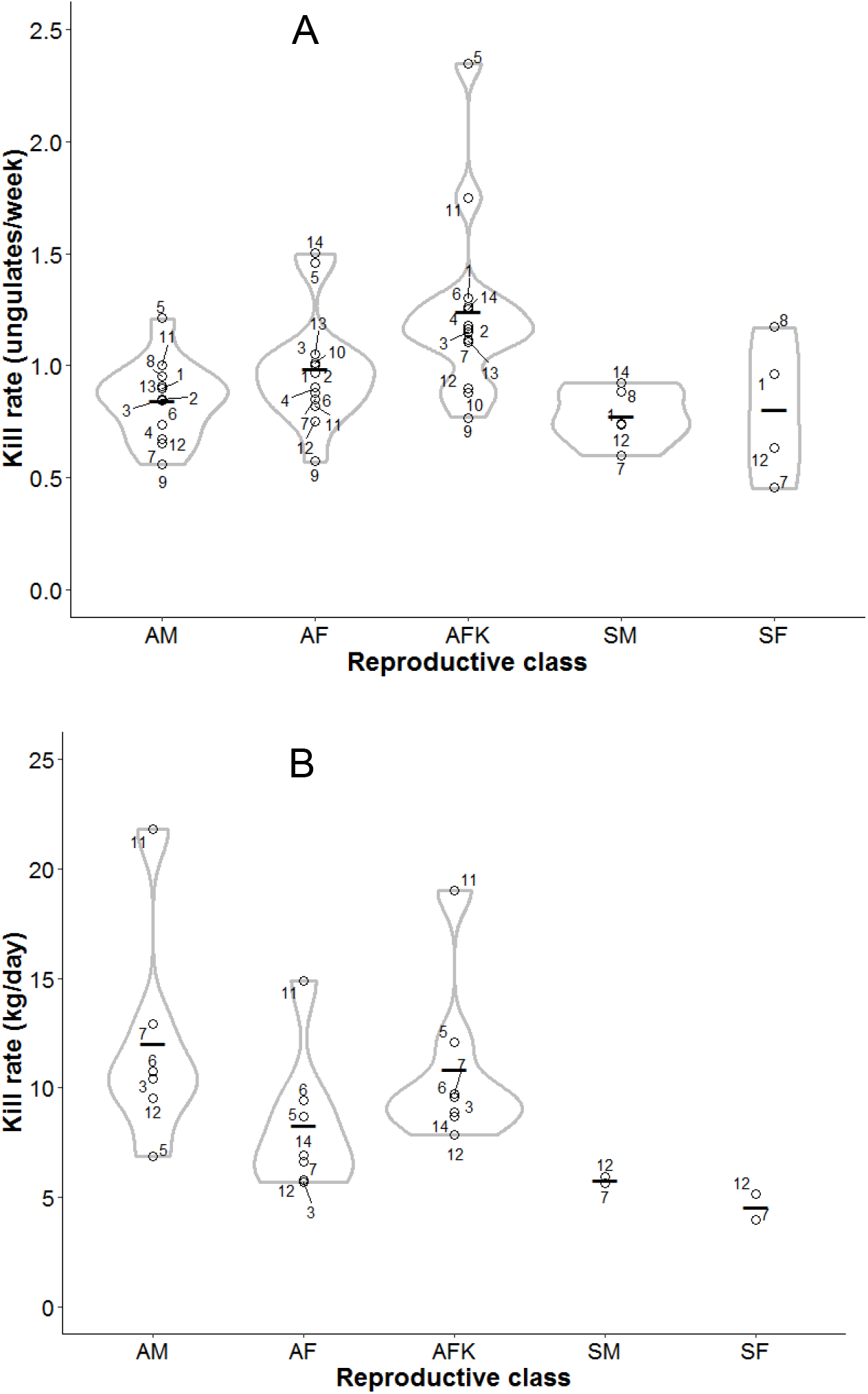
Puma (A) kill rate (ungulates/week) and (B) biomass killed (kg/day) across its distribution range. Horizontal black lines indicate mean kill rate. Gray contours show the distribution of the data, with widest areas denoting largest sample size of studies. Only studies that differentiated individuals by sex (male, female), age (adult, subadult) and adult females by reproductive status (solitary, with kitten(s)) are included. AM – adult male, AF – solitary adult female, AFK – adult female with kitten(s), SM – subadult male, SF – subadult female. Anderson & Lindzey 2003 (1), Blake and Gese 2016 (2), Clark et al. 2014 (3), Cooley et al. 2008 (4), Elbroch et al. 2014 (California) (5), Elbroch et al. 2014 (Colorado) (6), Knopff et al. 2010 (7), Mattson et al. 2007 (8), Mitchell 2013 (9), Nowak 1999 (10), Ruth et al. 2010 (11), Smith 2014 (12), White 2009 (13), Wilckens et al. 2016 (14).

The highest overall kill rate (2.35 ungulates/week) was reported for adult females with kittens in northern California (Elbroch et al. 2014) (Fig. 2A). This area had the second highest kill rate by solitary adult females (1.46 ungulates/week), a value only narrowly surpassed by frequency of kills made by this reproductive class in North Dakota (Wilckens et al. 2016). Northern California also had the only kill rate for adult males to exceed 1 ungulate/week (1.21 ungulates/week). Conversely, pumas inhabiting desert and xeric shrublands in Utah (Mitchell 2013) killed the least number of ungulates per week across adult reproductive classes (females with kittens 0.77, adult females 0.57, males 0.56).

#### Biomass

We found no significant differences in kill rates (kg/day) by adult reproductive class (*n* = 7) (Kruskal-Wallis 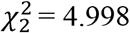, *P* = 0.082). However, on average adult males appeared to kill slightly more ungulate biomass (12.04 ± 5.17) than adult females with kittens (10.83 ± 3.84) and solitary females (8.28 ± 3.24) (Fig. 2B). Subadults were not included in the analysis due to small sample sizes, but males appeared to kill more biomass (5.77 ± 0.21) than females (4.56 ± 0.86). We also did not identify differences in biomass killed by reproductive class, when subsampling studies in Temperate Conifer Forests (*n* = 5) (Kruskal-Wallis 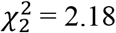, *P* = 0.336).

Pumas inhabiting the northern Yellowstone ecosystem consistently killed the largest ungulate biomass (kg/day) across all adult puma reproductive classes (males 21.80, solitary females 14.90, females with kittens 19.00; Fig. 2B) (Ruth et al. 2010). Minimum biomass killed per day was highly variable between study areas, but in all cases, adult pumas killed > 5 kg/day (males 6.86, solitary females 5.70, females with kittens 7.84).

### Factors Associated with Kill Rate

#### Frequency

Based on linear modelling, ungulate density was associated with puma kill rate for all adult puma reproductive classes (Table 3; *α* = 0.10). Pumas inhabiting study areas with high ungulate densities were more likely to have higher kill rates (Fig. 3), a result that was consistent for adult males (*F*_1,9_ = 5.08, *β* = 0.08 ± 0.04, *P* = 0.05), solitary adult females (*F*_1,10_ = 4.30, *β* = 0.09 ± 0.05, *P* = 0.06) and females with kitten(s) (*F*_1,10_ = 4.77, *β* = ± 0.09, *P* = 0.05). Kill rates of adult males were lower at high species richness of ungulates (*F*_1,9_ = 4.30, *β* = − 0.13 ± 0.06, *P* = 0.07). However, the data from California (Elbroch et al. 2014) was highly influential across the analyses (adult males: *D*_i_ = 1.20 and *D*_i_ = 0.81 > 0.44; solitary adult females: *D*_i_ = 2.36 > 0.40; females with kitten(s): *D*_i_ = 2.99 > 0.40). Subsetting the full dataset by excluding California and re-running the models resulted in loss of statistical support for these relationships. Overall, kill rate as a function of continuous covariates was best predicted by linear models. Generalized additive models were not supported except for adult male puma kill rate, for which a smooth term for ungulate density reduced to a linear term (Table 3, Supplementary Data S4).

**Table 3.**
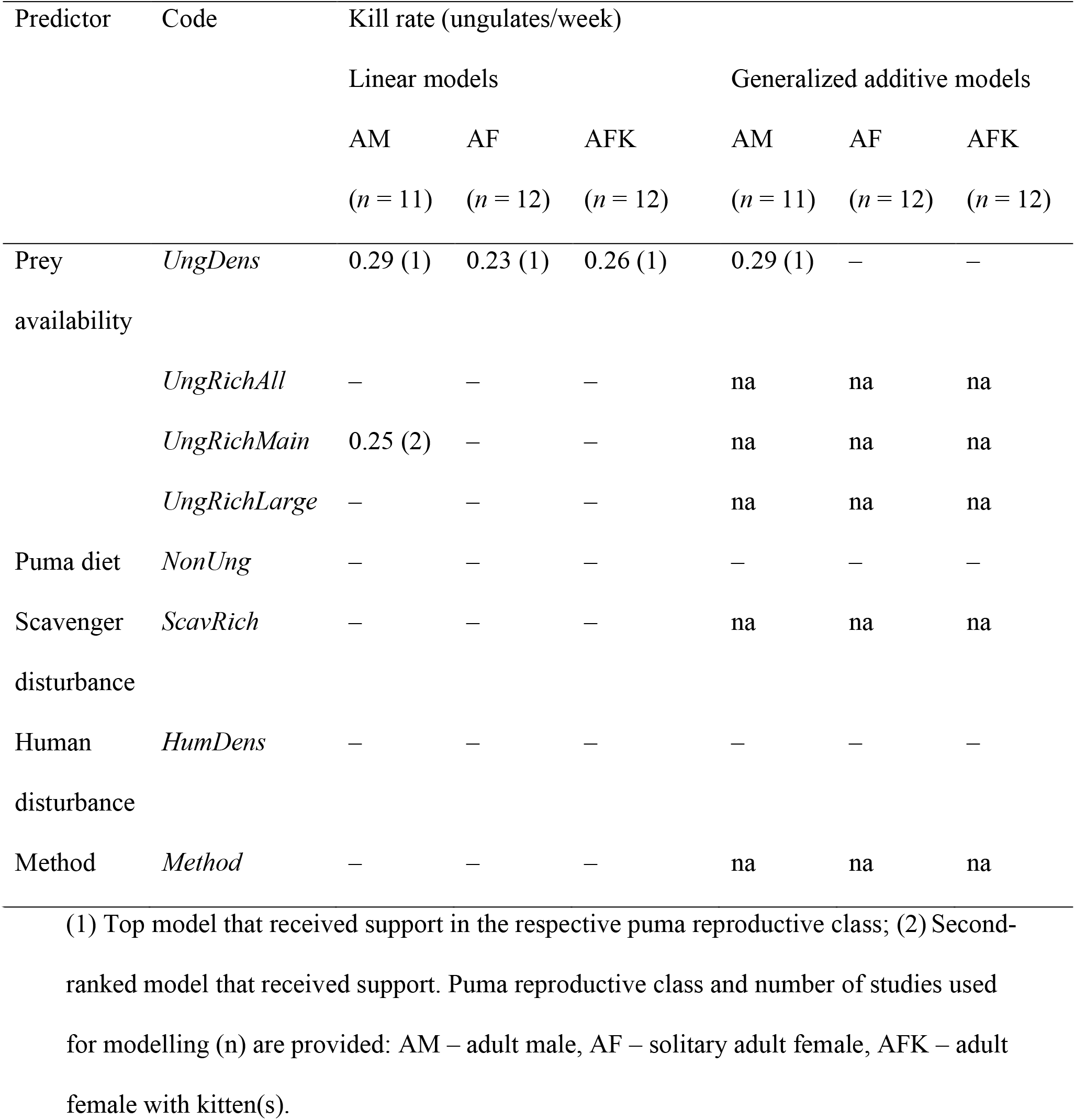
Support for predictors hypothesized to be associated with puma kill rate, based on studies throughout the puma distribution range. Values reported are the adjusted coefficient of determination (*R*^2^). Horizontal lines indicate no support for the respective models.

**Fig. 3.**
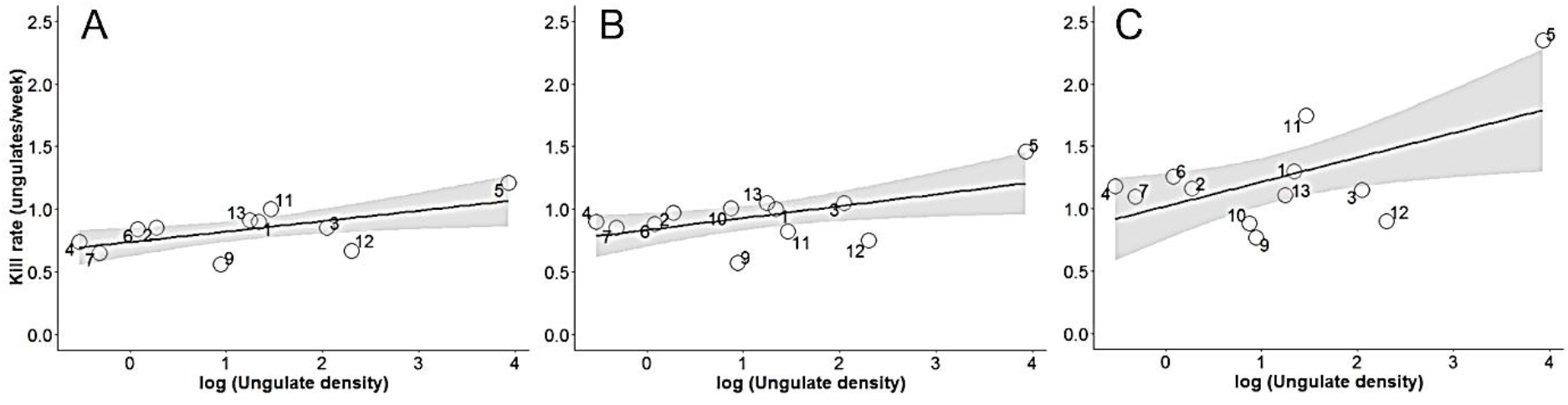
Relationship between puma kill rate (ungulates/week) and log ungulate density for (A) adult male pumas (*n* = 11), (B) solitary adult females (*n* = 12) and (C) adult females with kitten(s) (*n* = 12). Gray shaded areas are 90% confidence intervals. The sample sizes are the full datasets used for modelling kill rate as a function of factors potentially associated with it. Anderson & Lindzey 2003 (1), Blake and Gese 2016 (2), Clark et al. 2014 (3), Cooley et al. 2008 (4), Elbroch et al. 2014 (California) (5), Elbroch et al. 2014 (Colorado) (6), Knopff et al. 2010 (7), Mattson et al. 2007 (8), Mitchell 2013 (9), Nowak 1999 (10), Ruth et al. 2010 (11), Smith 2014 (12), White 2009 (13), Wilckens et al. 2016 (14).

#### Biomass

None of the models hypothesized to explain the biomass (kg/day) killed by pumas received support. However, this result should be interpreted with caution due to small sample (*n* = 6).

### Functional Response

Type I functional response curves had the best predictive ability for the datasets across all puma reproductive classes, suggesting that kill rates varied relatively linearly with ungulate density (Fig. 4). In contrast, Types II and III functional responses had larger and relatively similar mean predictive errors. In general, functional responses for adult males (*RMSE*_Type I_ = 0.125, *RMSE*_Type II_ = 0.153, *RMSE*_Type III_ = 0.159) had better fit than for adult females (*RMSE*_Type I_ = 0.138, *RMSE*_Type II_ = 0.197, *RMSE*_Type III_ = 0.202) and adult females with kittens (*RMSE*_Type I_ = 0.253, *RMSE*_Type II_ = 0.394, *RMSE*_Type III_ = 0.405).

**Fig. 4.**
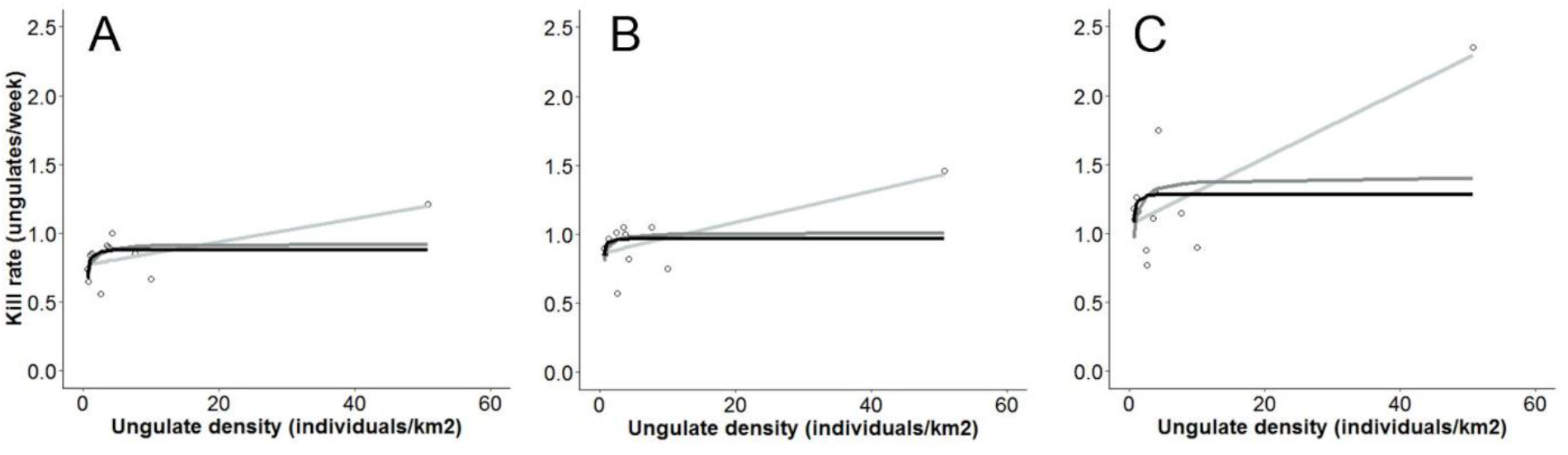
Functional responses of pumas across their range, including Type I (light gray), Type II (dark gray) and Type III (black). Data are for (A) adult male pumas (*n* = 11), (B) solitary adult females (*n* = 12) and (C) adult females with kitten(s) (*n* = 12).

## Discussion

Research on puma kill rates spans 70 years and 30 study systems across North and South America; it may be more extensive than similar research for any other apex predator, except perhaps gray wolves, and still our methods and inferences were severely hampered by the limited number of studies conducted to date and the inconsistency in kill rate study and reporting (Supplementary Data S1). Research on puma kill rates has disproportionately been conducted in Temperate Conifer Forests, and we could not find any studies from Tropical or Subtropical Moist Broadleaf Forests, which represent more than one third of puma range (Table 1). In North America, research is sparse for shrubland, desert and semi-desert systems. We found few kill rate studies of subadult pumas, and in general, young, dispersing pumas are the least studied age-class in terms of foraging ecology.

We only found six studies that reported kill rates in kg/day. The results of our analysis supported earlier work by Elbroch et al. (2014) and did not find differences in kill rates measured in kg/day across reproductive classes. Perhaps a larger sample size of studies might have yielded significant differences. Further, biomass metrics may provide distinctive insights into the energetic needs of top carnivores, as kill rate studies continue to accumulate, while accounting for ecological variables, such as scavenging. The importance of contrasting kill rates in kg/day and ungulates/week can be seen in data from the productive Yellowstone ecosystem: every adult puma reproductive class killed up to 3 times more in terms of biomass than in other systems for which we had comparable data (Ruth et al. 2010). Unlike many North American regions, elk are the main prey of pumas in the Greater Yellowstone Ecosystem (Elbroch et al. 2013; Ruth et al. 2019). In contrast, pumas in Yellowstone did not have the highest kill rates in terms of ungulates/week, suggesting a link between kill rate and prey size.

We discuss below the outcomes of testing predictors for kill rate frequency (ungulates/week), because the models hypothesized to explain biomass (kg/day) killed by pumas were not supported. We found support for prediction 1: Adult female pumas accompanied by kittens killed more ungulates per unit time than the other reproductive classes. This pattern has been documented for other solitary carnivores killing primarily wild ungulates (e.g. Eurasian lynx, *Lynx lynx*, Andrén and Liberg 2015; cheetah, *Acinonyx jubatus*, Hilborn 2017), which is likely driven by increased energetic requirements of family groups as compared to solitary individuals (Laundré 2005). Despite weighing substantially more than adult females, the energetic needs of adult males are likely lower than for females with kittens. Adult males also kill larger ungulates than adult females in some systems (Knopff et al. 2010; Elbroch et al. 2013; Clark et al. 2014), and the greater biomass of larger prey may reduce their kill rates. Adult males may also be more effective than females at defending carcasses against scavengers, as competition is largely dictated by the size of competitors (Donadio and Buskirk 2006). Finally, males feed from kills made by females within their territories in some systems, which may reduce their kill rates, while the reverse appears much less common (Elbroch et al. 2017b). Subadult data were insufficient for analyses, but raw values of subadult kill rates were lower than for adults, matching recent research showing that subadults consume disproportionately more small-bodied, non-ungulate prey (Elbroch et al. 2017a), and that pumas prey switch as they age and refine their hunting skills (Elbroch and Quigley 2019).

We found partial support for prediction 2, that prey availability is positively associated with puma kill rate. Ungulate density was the most important factor consistently associated with kill rate across adult puma reproductive classes. Analysis showed higher kill rates at higher ungulate density, in accordance with a Type I functional response. The adult male Type I functional response had the best fit compared to adult females, irrespective of whether the latter were solitary or accompanied by kittens. Biologically, this might possibly suggest that males may be able to kill ad libitum, which would then allow them to spend more time fulfilling other life history activities, such as territorial marking and searching for females. Alternatively, our results may indicate that we did not detect the threshold at which functional responses asymptote, as in Type II and III functional responses (Holling 1959). These findings, however, must be interpreted with caution, for prey size likely explained some variation across study sites, as exemplified above in Yellowstone. Influential observations also had a large effect on findings. Overall, the highest kill rates (ungulates/week) were recorded for California’s North Coast. This productive ecosystem had disproportionately higher ungulate density than any other area analyzed (50.75 ungulates/km^2^) (Lounsberry et al. 2015). When we removed this one data point (California; Elbroch et al. 2014), ungulate density no longer correlated with kill rates for any adult reproductive class. Similarly, when California data was removed, ungulate richness no longer correlated with and explained adult male kill rates.

We did not find support for prediction 3, that puma populations with a high proportion of non-ungulate prey in their diet would have lower ungulate kill rates. This could be due to logistics, as some small non-ungulate prey might go undetected during field visitation of GPS collar location clusters (Bacon et al. 2011). Or it could be because of the disproportionate energetic value of ungulates, which often weigh ≥ 10x non-ungulate prey (e.g. Knopff et al. 2010). In addition, the agencies that typically fund this work may deemphasize research focus on secondary prey, and thus field staff may not prioritize investigating GPS clusters of short duration. Although this aspect of puma foraging ecology may appear less relevant to wildlife managers, abundant secondary prey could impact puma-primary prey interactions. For example, abundant secondary prey appear to be hunted opportunistically (Cristescu et al. 2019) and therefore are likely included in puma diet as explained by their abundance. Increased foraging resources may also influence puma recruitment, especially survival of subadults if young animals select for non-ungulate prey during dispersal (Elbroch et al. 2017a). Thus, better understanding puma utilization of non-ungulate prey has implications for the management of ungulates as well.

We did not find support for prediction 4, that kill rates would positively correlate with dominant scavenger richness, but we interpret these results with caution. Dominant scavenger abundance would have been a better covariate to test against puma kill rate, but we lacked such data. Alternatively, pumas may exhibit high kill rates that buffer against the effects of dominant competitors because they evolved to withstand high levels of kleptoparasitism (Elbroch et al. 2017c), as has also been suggested for cheetahs (Scantlebury et al. 2014). Further research is needed to determine threshold scavenger impacts on puma fitness.

Neither did we find support for prediction 5, that kill rates would positively correlate with human density, but as with scavengers, this may have been due to our methods rather than reflective of ecological systems. Human density influenced kill rate in an urban setting (Smith et al. 2015), but few studies in our analysis occurred in such areas. In addition, we calculated human density at the level of a study area, whereas Smith et al. (2015) were able to quantify this variable at the level of a puma home range. We suggest that future studies, especially those in urbanized areas, collect and include data on the intensity of recreation—at the home range level if possible—rather than strict human census data, as this may be more informative in determining the impact of human activity on puma behaviors (*sensu* Smith et al. 2017).

Surprisingly, we did not find support for prediction 6, that field methodology (VHF vs. GPS) would impact kill rate values. This may have been due to our decision to include studies that estimated kill rate after visiting a subset of kills in the field and thereafter employing predictive modeling to assess the probability of kills made by pumas over time. Predictive modeling has been shown to underestimate kill rates in some systems (Elbroch et al. 2018), and therefore their inclusion may have diluted the differences between VHF and GPS studies. Ruth et al. (2010) reported similar results at the scale of one study area, but they visited a subset of GPS clusters and used predictive modeling to estimate kill rates.

In reality, all of our predicted different influences on puma kill rates may be additive or synergistic, complicating ecological inferences. For example, we documented the highest kill rate in ungulates/week in northern California, which exhibited the highest ungulate density but the smallest ungulate prey species, black-tailed deer (*O. h. columbianus*). Further, the California system included high densities of American black bears, which displaced pumas from their kills over most of the year, because it is a warmer climate where bears hibernate for a shorter duration than in other parts of their range (Allen et al. 2014; Allen et al. 2015). In Yellowstone, where kill rates in kg/day were the highest (Ruth et al. 2010) and pumas primarily predated elk (Ruth et al. 2019), the scavenger community is the most complex, with American black bears, grizzly bears and gray wolves displacing pumas from their kills. In systems where pumas are frequently displaced by scavengers or people, they may need to increase kill rates to compensate for losses (Elbroch et al. 2015b; Smith et al. 2015). In contrast, the lowest kill rates (ungulates/week) were recorded for pumas in a semi-arid region in Utah (Mitchell 2013). This system had a considerably lower ungulate density than California’s North Coast (2.56 ungulates/km^2^) and lacked dominant scavengers completely, which may have allowed pumas to consume more of their kills. In addition, the low humidity of arid regions desiccates fly eggs and affects the growth of larval stages on carrion (Forbes and Carter 2015), possibly extending handling times which may lower puma kill rates in these regions. Lastly, carnivore personality and intraspecific variation in prey selection could also complicate inferences (Pettorelli et al. 2011), especially at low sample sizes of individuals monitored, as is common for projects in which researchers estimate kill rates.

### Conclusions

Although traditionally used to infer the effects of carnivores on ungulate populations, kill rates are most useful to advance our understanding of the foraging ecology of predators. Kill rate studies by themselves do not provide much insight into prey population dynamics (Vucetich et al. 2011), unless augmented with predator and prey density, prey survival and fecundity data, and ideally, data quantifying the additional effects of competitors or scavengers. In our review, kill rates were highest for adult females with kittens, suggesting that an abundant prey base is necessary to sustain this reproductive class, which is an important consideration for endangered or threatened carnivores. Kill rates measured in biomass, on the other hand, varied to a lesser extent among reproductive classes. We found evidence that kill rates were higher at high prey density, a relationship that was mostly driven by one single study area and should therefore be interpreted with caution. The large influence of small sample sizes highlighted the need for further research on kill rates of pumas and other top carnivores across biomes, as well as replication within biomes, to allow for a more robust assessment of ecological and behavioral factors driving kill rates and functional response. Nonetheless, the positive association between prey density and kill rate illustrates the value of inference across multiple studies for investigating the shape of the functional response. Such empirical assessments of the relationship between predators and prey are generally not possible to obtain from one area alone, and have rarely been implemented across study systems involving large carnivores.

With additional research across the distribution of apex predators, it will be possible to perform multivariate analyses, and interact variables of interest at continental and global scales, as is a growing trend in ecological research (Schimel and Keller 2015; Steenweg et al. 2017). To maximize the benefits of future studies requires standardizing research methods and reporting in kill rate studies:

1. The methodology for estimating kill rates should involve GPS collars and significant time in the field differentiating kill sites from sites associated with other activities (Elbroch et al. 2018), preferably in conjunction with accelerometer devices that can greatly improve the accuracy of identifying both kill and scavenging events (Wang et al. 2015; Petroelje et al. 2020). GPS clusters are well suited for locating predation events of large and medium carnivores (Merrill et al. 2010; Jansen et al. 2019).
2. Kill rate sampling should take place across seasons. For example, in the northern hemisphere, summer kill rates are generally higher than winter kill rates, because many carnivores select for newborn ungulates at this time of year (Knopff et al. 2010; Allen et al. 2014), and perhaps also due to the seasonal activities of dominant scavengers such as bears (Elbroch et al. 2014).
3. Kill rates should be reported in ungulates per unit time, that is all species of ungulates in multi-prey systems, and in kg/day. Kill rates can also be reported for specific ungulates, but this should complement rather than replace kill rates measured for all ungulates. Where small prey contributes substantially to carnivore diet, kill rates could additionally be reported for small prey per unit time.
4. All diet items, and their sex and age class, should be listed in supplementary materials, in order to allow others to recreate kill rates in kg/day or other biomass metrics.

## Acknowledgments

We thank the California Department of Fish and Wildlife for funding support. We are grateful to J. Heiman, J. Kolek, E. Leonhardt, R. Lyon and E. Wildey, for assistance with literature searches.

## Supplementary Data

Supplementary Data S1. Kill rate studies identified in the review of puma kill rate across its biogeographical range.

Supplementary Data S2. Values of the predictor variables used in modelling puma kill rate (ungulates/week) across its biogeographical range.

Supplementary Data S3. Values of the predictor variables used in modelling puma kill rate (kg/day) across its biogeographical range.

Supplementary Data S4. Plot of predicted values of adult male puma kill rate based on a generalized additive model with a single smooth term for ungulate density.

