## Supplementary figures and images for "Biogeographical and ecological factors associated with kill rates of an apex predator"

### Supplementary Data S4

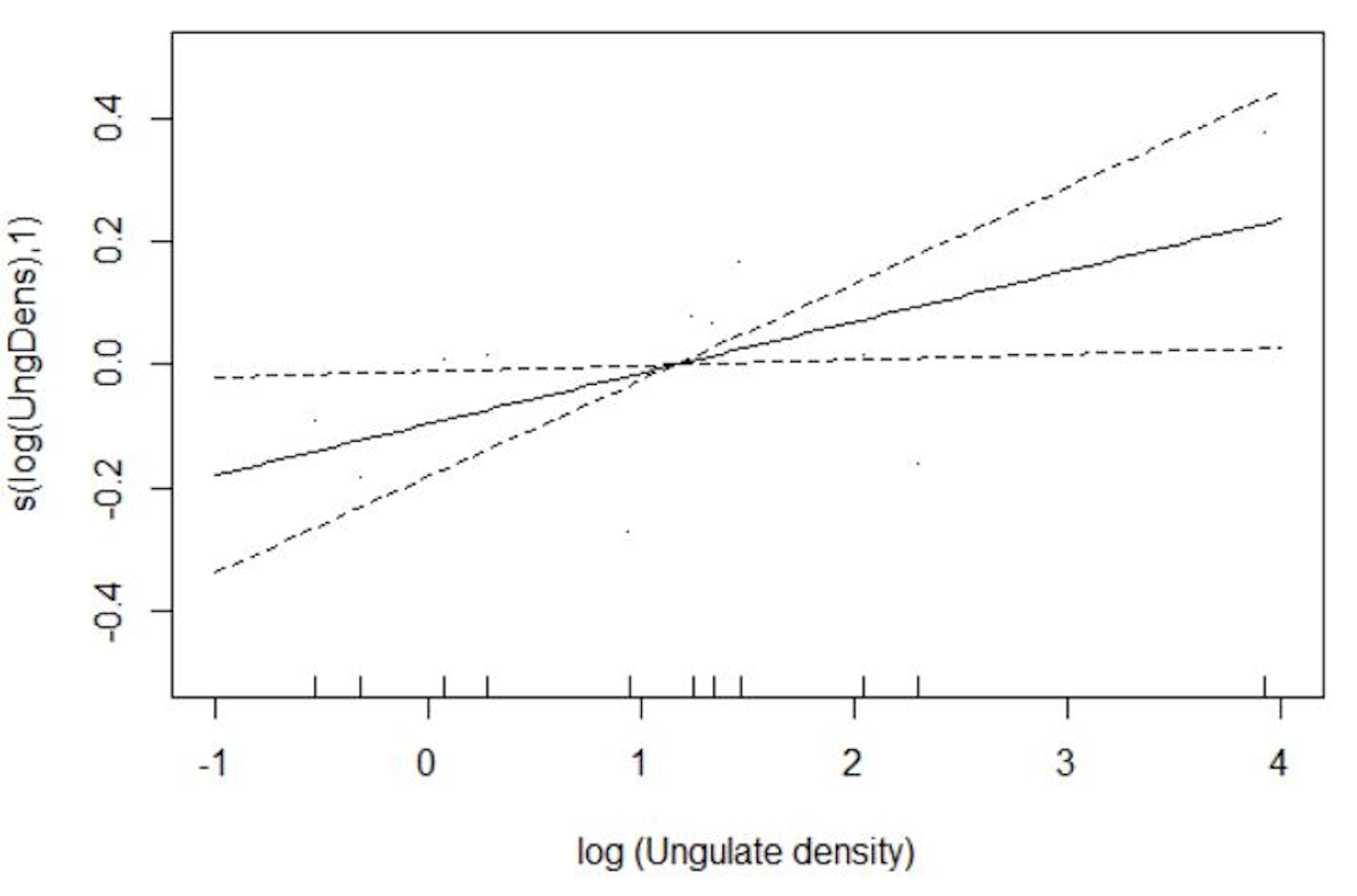
